# Wide field-of-view, multi-region two-photon imaging of neuronal activity *in vivo*

**DOI:** 10.1101/011320

**Authors:** Jeffrey N. Stirman, Ikuko T. Smith, Michael W. Kudenov, Spencer L. Smith

## Abstract

We demonstrate a two-photon imaging system with corrected optics including a custom objective that provides cellular resolution across a 3.5 mm field of view (9.6 mm^2^). Temporally multiplexed excitation pathways can be independently repositioned in XY and Z to simultaneously image regions within the expanded field of view. We used this new imaging system to measure activity correlations between neurons in different cortical areas in awake mice.

---

Two-photon^1^ population calcium imaging *in vivo*^2^ supports dense optical sampling of neuronal activity deep in scattering tissue such as mammalian neocortex and unambiguous identification of recorded neurons, particularly when genetically encoded indicators are employed^3,4^. This approach is used to measure single cell-level stimulus selectivity in dense local ensembles of neurons^5,6^ and learning and task-related activity in awake mice^7^, including those navigating in a virtual reality environment^8^. Conventional two-photon imaging systems offer high resolution across a field of view (FOV) that is typically smaller than a cortical area in a mouse (*e.g.,* primary visual cortex). However, neuronal activity that supports sensory coding and motor output is distributed across multiple areas and millimeters of neocortex^9,10^ (**Fig. 1a**), thus a cellular-resolution view into ongoing neural activity across extended cortical networks is essential for elucidating principles of neural coding.

**Figure 1.**
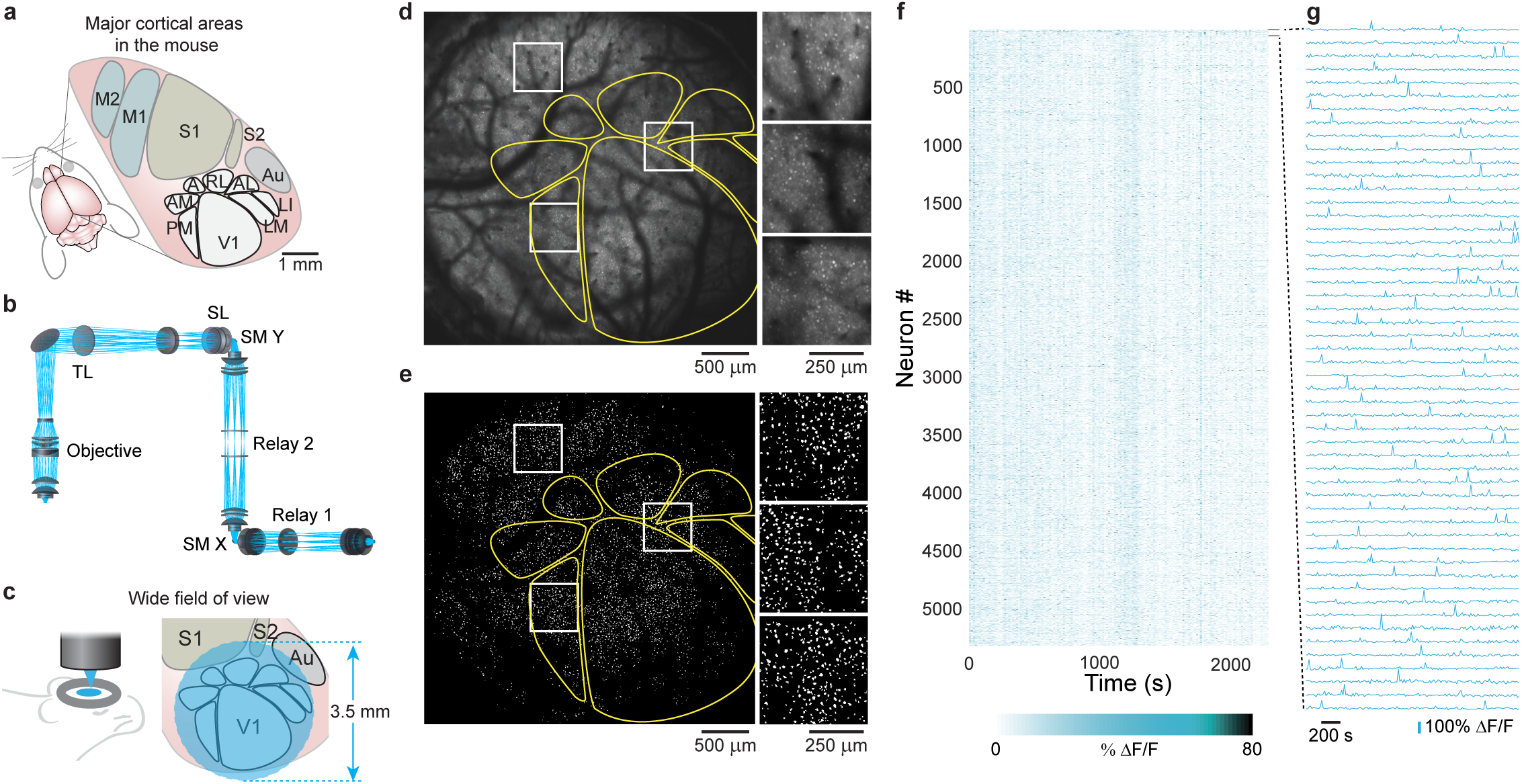
Two-photon imaging of neural activity across 9.6 mm^2^ of mouse cortex with single neuron resolution. (**a**) In the mouse, primary visual cortex (V1) is surrounded by higher visual areas (HVAs; PM = posteromedial; AM = anteromedial; A = anterior; RL = rostrolateral; AL = anterolateral; LM = lateromedial; LI = laterointermediate), which are distributed across several millimeters of cortex (M1,M2 = primary and secondary motor cortex; S1,S2 = primary and secondary somatosensory cortex; Au = auditory cortex). A wide field of view (FOV) is required to image neuronal activity in these distributed cortical areas simultaneously. (**b**) Multi-lens optical systems were developed to realize a laser scanning 2p microscope system with a large FOV while still preserving single neuron resolution (TL = tube lens; SL = scan lens; SM X, SM Y = scan mirrors for X and Y scanning). (**c**) The 3.5 mm FOV can encompass V1 and HVAs. (**d**) A transgenic mouse expressing the genetically encoded fluorescent calcium indicator GCaMP6s in excitatory neurons was used to examine neuronal activity (maximum projection). Prior to 2p imaging, intrinsic signal optical imaging was used to map out the higher visual areas (yellow outlines). The expanded inlays (white) show cellular resolution is preserved across the FOV (also see **Supplementary Figure 9** and **Supplementary Video 3**). (**e**) Segmenting the image sequence yields 5,361 active neurons, (**f**) whose visually-evoked responses were recorded, and (**g**) revealing clear fluorescence transients (traces for the first 50 neurons in panel **f**).

Here we present a two-photon imaging system with custom optics that preserves single neuron resolution across a wide FOV (3.5 mm diameter; 9.6 mm^2^). To simultaneously image neurons over extended cortical networks, dynamically and independently repositionable temporally multiplexed imaging beams are synchronously scanned. We refer to the system as a Trepan2p (<u>T</u>win <u>R</u>egion, <u>Pan</u>oramic <u>2</u>-<u>p</u>hoton) microscope.

Multi-element optical subsystems (afocal relays, scan lens, tube lens, and objective) were designed (**Fig. 1b**, **, Supplementary Figs. 1-4, Online Methods**) to minimize aberrations at the excitation wavelength (910 ± 10 nm) across scan angles up to ± 4 degrees at the objective (**Supplementary Fig. 5a**). The development of custom optics was necessary because commercially available microscopes do not provide objectives and scan optics that support 2p imaging with single neuron resolution across a large FOV. Commercially available low magnification objectives lack sufficient axial resolution and collection efficiency to image individual neurons at depth *in vivo* (though in a recent study, a custom scan engine combined with a commercial objective was used to image blood flow^11^). Large diameter macroscope objectives that offer a wide field of view are commercially available, but they and their associated scanning optics are not optimized for multiphoton excitation, which is acutely sensitive to optical aberrations and requires large beam diameters and high scan angles to use the full NA and FOV. Most commercial objectives are well corrected on axis and off axis up to a few (2-3 degrees), but their performance dramatically drops off significantly at larger angles. Furthermore, many commercial scanning systems (relay, scan lens, tube lens) can exhibit clipping at scan angles over 1-2 degrees, and may not be well corrected for aberrations at those angles. Either problem can degrade (expand) the point spread function (PSF). In two-photon imaging, expanding PSFs can geometrically decrease the excitation efficiency and thus limit the FOV due to vignetting (dark areas outside of a central region).

In designing the custom optics, we prioritized the constant performance (RMS wavefront error) over the entire designed scanning range. In this way we could preserve cellular resolution anywhere within the large FOV. Subsystems can be diffraction limited on their own, but demonstrate additive aberrations when used together in a full imaging system. Optimizing the system as a whole (including all relays, scan lens, tube lens, and the objective), rather than optimizing components individually, ensured we would meet the desired performance. The custom objective (27.5 mm effective focal length, 0.4 NA), together with the scan optics that were highly corrected to minimize aberrations, yielded single neuron resolution with two-photon excitation across a 3.5 mm FOV (**Fig. 1c**). We evaluated the resolution of the Trepan2p microscope by measuring the excitation volume as the full-width at half-maximum (FWHM) of the intensity profile of 0.2 μm beads (**Online Methods**). Radial FWHM was ∼1.2 ± 0.1 μm (mean ± SD) both at the center and at the edges of the FOV (1.75 mm from the center). The axial FWHM was 12.1 ± 0.3 μm at the center, and 11.8 ± 0.4 μm at the edges of the FOV (both measurements are mean ± SD; **Supplementary Fig. 6**). This resolution is sufficient to image activity with single neuron resolution in mouse neocortex (**Supplementary Video 1, Supplementary Note 1**)^12^ as well as resolve sub-cellular structures such as dendritic spines (**Supplementary Fig. 7**). Furthermore, the moderate NA of the objective can aid imaging deep in scattering tissue, compared to objectives with NAs > 1.0 (refs. ^13,14^). We were able to image the somas of Layer 5 neurons over 700 μm below the cortical surface *in vivo* (**Supplementary Video 2**). Field curvature and distortion were minor (**Supplementary Fig. 5c, Online Methods**). Thus, across the full FOV, cellular resolution was maintained, and for a sampling interval of 0.5 μm, the full FOV contained 38.5 megapixels.

To demonstrate the functionality of the Trepan2p, we performed two sets of *in vivo* imaging in transgenic mice that expressed the genetically encoded calcium indicator, GCaMP6s^3^, in neocortical pyramidal neurons. In the first set of experiments, we explored the optical access to neurons across the large FOV. To locate primary visual cortex (V1) and higher visual areas (HVAs) in the mouse^10^, we used intrinsic signal optical imaging^15^ (**Online Methods, Supplementary Fig. 8**). We then imaged neuronal activity-related GCaMP6 signals in a large cortical region (**Fig, 1d, Supplementary Fig. 9**, **Supplementary Video 3**). The full FOV was scanned with a single beam at ∼0.1 frame/s (3.5 mm wide FOV; 2048 × 2048 pixels; 3 μs dwell time per pixel), and contained in it V1 and at least six HVAs. Undersampling and alternative scan strategies such as <u>arbitrary line scanning</u> (**Supplementary Fig. 10**) can decrease the acquisition time per frame^16,17^, <u>and stimulus presentation durations can facilitate the capture of stimulus-related events</u>. Neurons were readily detected throughout the entire FOV based on spiking activity reported by GCaMP6s (5361 neurons detected, **Online Methods; Fig. 1e-g**). Thus the Trepan2p system provides cellular-resolution optical access across the full FOV (9.6 mm^2^), which encompasses more than six different mouse cortical areas within a single FOV.

Investigations of neural correlations and dynamics can require simultaneously measuring activity in two different cortical areas. To simultaneously scan two regions within the full FOV at rates fast enough to image activity dynamics during behavior, we developed a repositionable beam system with two temporally multiplexed imaging pathways^18-20^(**Fig. 2a, Supplementary Fig. 11a**). Custom motorized steering mirrors (SM1 and SM2), conjugated to the scan mirrors using afocal relays, impart independent solid angle deflections (*Ω*_1_ or *Ω*_2_) for the two beams and the <u>focal plane for each pathway is adjusted</u> using a tunable lens^21^ (**Fig. 2a**). Thus, each path is independently positionable in *X, Y,* and *Z* at will during imaging without moving the microscope or the preparation (**Fig. 2b**). As the rest of the system, the beam recombination optics were designed for low aberrations to ensure effective 2p excitation and resolution across the wide FOV. The single photon PMT pulses are demultiplexed using a synchronization signal from the laser module, assigning detected photons to pixels in the associated imaging pathway with minimal crosstalk between pathways (**Online Methods, Supplementary Figs. 11b,c, 12**).

**Figure 2.**
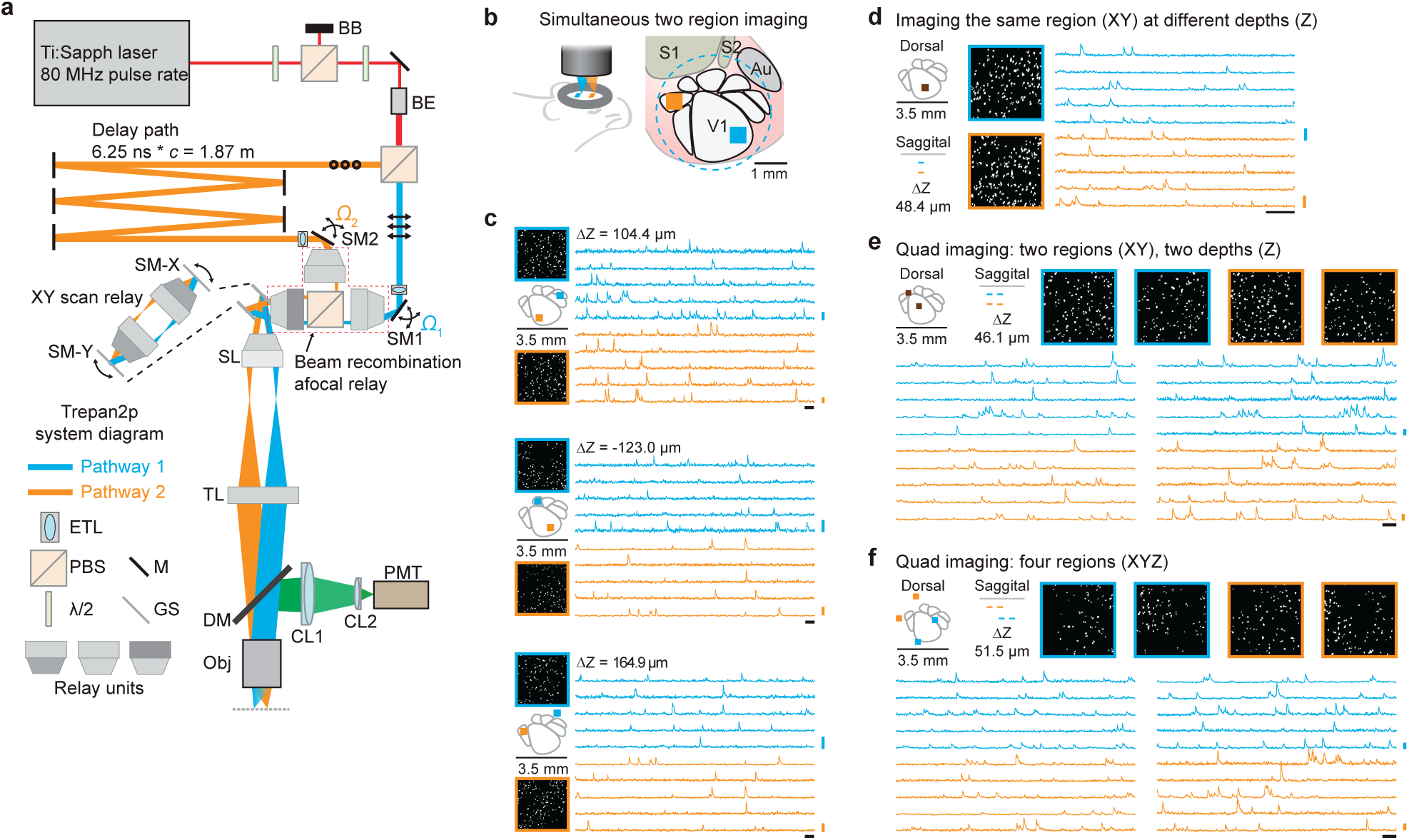
Temporally multiplexed, independently repositionable imaging pathways for simultaneous scanning two regions. (**a**) Two imaging beams are temporally multiplexed and independently positioned in XY and Z prior to the scan mirrors (SM-X, SM-Y). First, overall power is attenuated using a half-wave plate (λ/2), a polarizing beam splitting cube (PBS) and a beam block (BB). After a second λ/2 (used to determine the power ratio sent to the two pathways) and a beam expander (BE), a second PBS divides the beam into two pathways. Pathway 1 (in blue, p-polarization, indicated by the arrows) passes directly to a motorized steering mirror (SM1) for positioning in *XY*. Pathway 2 (in orange, s-polarization, indicated by the circles) passes to a delay arm where it travels 1.87 meters further than pathway 1 using mirrors (M), thus delaying it by 6.25 ns before being directed to SM2. Directly prior to SM1 and SM2 are electrically tunable lenses (ETL) that can adjust the *Z* position (focal plane) of the pathways independently. The two pathways are recombined (beam recombination relay), and sent to X and Y galvanometer scanners (GS) that are connected by an afocal relay (expanded view inset). A scan lens (SL) and tube lens (TL) focuses the two multiplexed beams onto the back aperture of the objective (Obj). Fluorescence is directed to a photomultiplier tube (PMT) via an infrared-passing dichroic mirror (DM) and two collection lenses (CL1, CL2). (**b**) The individual imaging regions can be independently positioned and repositioned anywhere within the full FOV by the steering mirrors (SM1,SM2 in **a**) for *XY* position, and the tunable lenses (ETL in **a**) for independent *Z* positioning. (**c**) Within the same session, without moving the mouse or the microscope, the two pathways were moved to various configurations (left) to image neuronal activity (3.8 frames/s) (left, segmented active ROIs within the 500 μm imaging region; right, five example traces from each region). Relative axial displacement (ΔZ) of the focal planes of the two imaging pathways compensated for brain curvature. (**d**) There is no lower limit to the *XYZ* separation between imaging pathways. In this imaging session (9.5 frames/s), the *XY* locations were identical and the pathways only differed in the *Z* depth (left, segmented active ROIs within the 500 μm imaging region; right, five example traces from each region; **Supplementary Video 4**). (**e, f**) By combing temporal multiplexing (Pathways 1 and 2) with serially changing the offset voltage on the galvanometer scanner, four regions can be rapidly imaged (10 frames/s). (**e**) Pathway 1 and 2 are positioned at the same XY location and offset in Z. The galvanometers serially position the imaging region (of each pathway) anywhere within the larger field of view (left, segmented active ROIs within the 400 μm imaging region; right, five example traces from each region; **Supplementary Video 5, 6**) (**f**) Pathway 1 and 2 are positioned at different XY locations as well as offset in Z. The galvanometers serially position the imaging region (of each pathway) anywhere within the larger field of view (left, segmented active ROIs within the 400 μm imaging region; right, five example traces from each region). For all panels the vertical scale bar is 200% ΔF/F and the horizontal scale bar is 10 s.

To demonstrate the flexibility of this approach, we performed experiments with a variety of configurations of the twin imaging pathways. While presenting visual stimuli to the mouse (a naturalistic movie), we simultaneously recorded activity in several different pairs of cortical areas, and at different depths (512 × 512 pixels, 3.8 frames/s, **Fig. 2c)**. The imaging regions can be placed arbitrarily close to each other. To demonstrate this, we placed the two imaging paths at the same *XY* location, but offset their positions in *Z* (512 × 256 pixels, 9.5 frames/s, **Fig. 2d, Supplementary Video 4**). Next we explored quad region imaging. Each of the dual imaging pathways were configured to alternate (every other frame) between two different regions. Thus, we imaged four different XY regions, each at 10 frames/s (250 × 100 pixels, **Fig. 2e**; 512 × 512 pixels, 3.8 frames/s, **Supplementary Videos 5, 6**). We also imaged four regions that differed in the XY locations and Z (250 × 100 pixels, 10 frames/s, **Fig. 2f**).

In another combination, we imaged in both V1 and in medial higher visual areas (HVAs) AM and PM (<u>250 × 100 pixels, 20 frames/s</u>, **Fig. 3a**) while presenting a pair of visual stimuli (a naturalistic movie or drifting gratings). Pairwise correlations between cortical areas were calculated from spike time courses inferred from the GCaMP6s signals (**Supplementary Note 2**). This analysis revealed an increase in correlated activity between neurons in V1 and neurons in areas AM and PM when the naturalistic movie is presented, compared to gratings (cross-correlation with gratings (mean ± SEM): 0.0157 ± 0.0003; with naturalistic movie: 0.0218 ± 0.0003; *N* = 12160 neuron pairs; *P* < 10^-10^; rank-sum test; **Fig. 3b**). Such cross-area correlation analysis between densely sampled populations of neurons was made possible with the large FOV and temporally multiplexed scanning. These imaging regions can be as large or small as needed for the experiment (for example, a 2.5 mm FOV at two different z-depths; 2048 × 1024 pixels; 0.8 frames/s); **Supplementary Fig. 13; Supplementary Video 7**), and two regions can be set to scan complementary halves of the FOV (as in ref. ^18^). Thus, the repositionable temporally multiplexed imaging beams can be reconfigured within an experiment for a variety of measurements.

**Figure 3.**
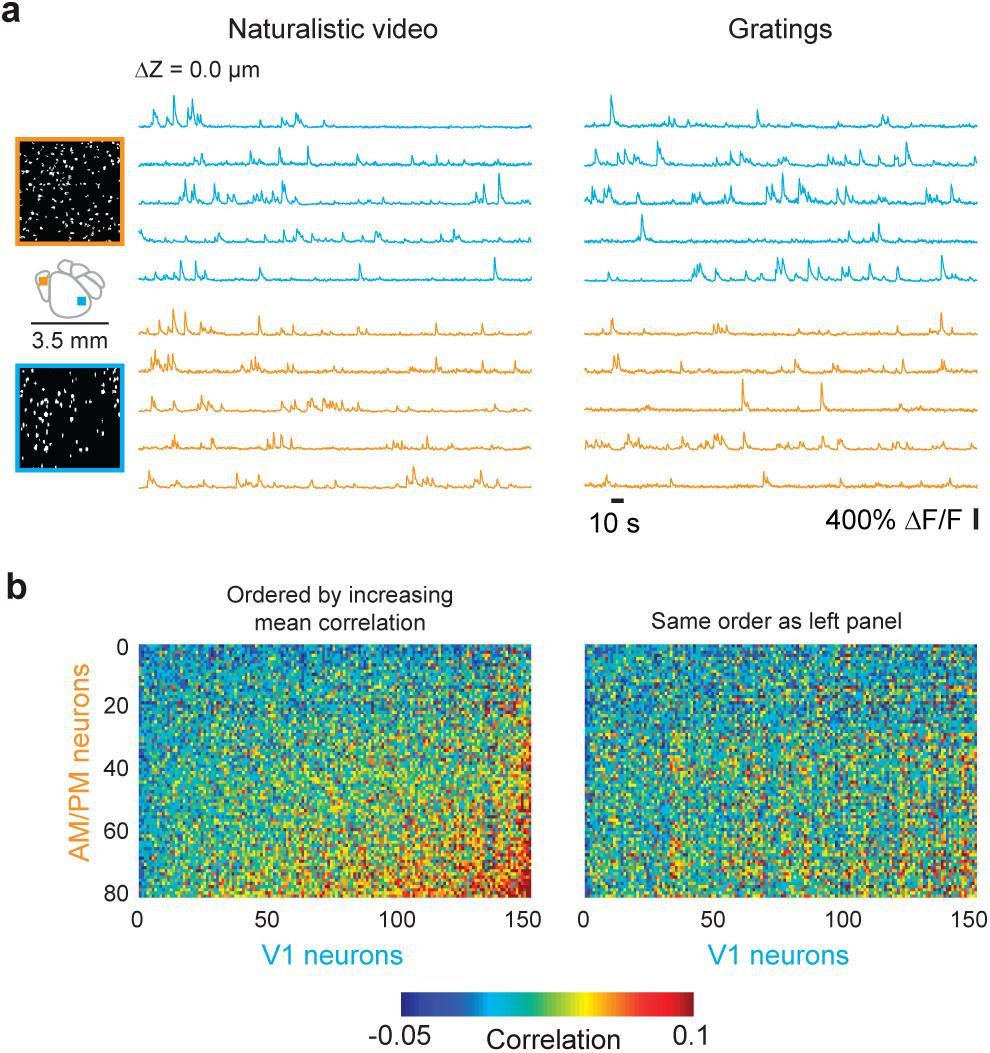
Simultaneous imaging in V1 and AM/PM to explore stimulus dependent changes in correlation structure. (**a**) Neuronal activity was imaged in two regions (20 frames/s), V1 and in an ROI encompassing retinotopically matched regions of AM and PM, simultaneously. Visual stimuli, either drifting gratings or a naturalistic movie, were used to evoke responses (left, segmented active ROIs within the 400 μm imaging region; right, five example traces from each region and for each visual stimulus). (**b**) Ca^2+^ signals were used to infer spike times and examine correlations. Activity correlations were measured between pairs of cells, each pair consisting of a V1 neuron and a neuron in AM or PM. These correlations were higher during presentation of the naturalistic movie compared to those during the drifting gratings (cross correlation with gratings, mean ± SEM: 0.0157 ± 0.0003; with naturalistic movie: 0.0218 ± 0.0003; *N* = 12160 neuron pairs; *P* < 10^-10^; rank-sum test). The neurons on both axes were ordered from low to high average correlation for presentation clarity on the left (naturalistic movie), and the same ordering is used on the right (gratings).

These results demonstrate the utility of the Trepan2p system. The large working distance (8 mm) of the air-immersion objective can facilitate integration with head-fixed behavior experiments. Moreover, in principle the same instrumentation can be applied to image neural activity in many animal models whose cortical organization extends beyond the spatial limits of conventional 2p microscopy, such as multiple orientation columns in primary visual cortex of cats or non-human primates. The methodological approach presented here provides a flexible platform for measuring neural activity correlations and dynamics across extended cortical circuitry with single neuron resolution.

## METHODS

Methods and any associated references are available in the online version of the paper.

## ACKNOWLEDGEMENTS

We are grateful to Hongkui Zeng for kindly providing transgenic mice, Kei Eto for providing mice for imaging in pilot experiments, Janet Berrios for providing the YFP labeled brain section, Sally Kim and Ben Philpot for providing the Thy1-GFP O-line mouse, and Zemax LLC for providing upgraded software. This work was supported by the following: the National Institute of Child Health and Human Development (T32-HD40127) and Burroughs Wellcome Fund Career Award at the Scientific Interface (J.N.S.); a Helen Lyng White Fellowship (I.T.S.); and a Career Development Award from the Human Frontier Science Program (00063/2012), and grants from the National Science Foundation (1450824) and the Whitehall Foundation (S.L.S.).

## AUTHOR CONTRIBUTIONS

S.L.S. conceived the Trepan2p imaging system. J.N.S. and S.L.S. designed and engineered the system. J.N.S. developed the demultiplexing electronics, optimized the optical systems, wrote the software, and constructed the system. M.W.K. and J.N.S. modeled optical subsystems and M.W.K. consulted on optical optimizations. J.N.S., I.T.S., and S.L.S. performed the animal experiments. J.N.S. and S.L.S. analyzed and interpreted the data. J.N.S. and S.L.S. wrote the manuscript with input from all authors. I.T.S. and S.L.S. supervised the project.

## Supplementary Figures

**Supplementary Figure 1.**
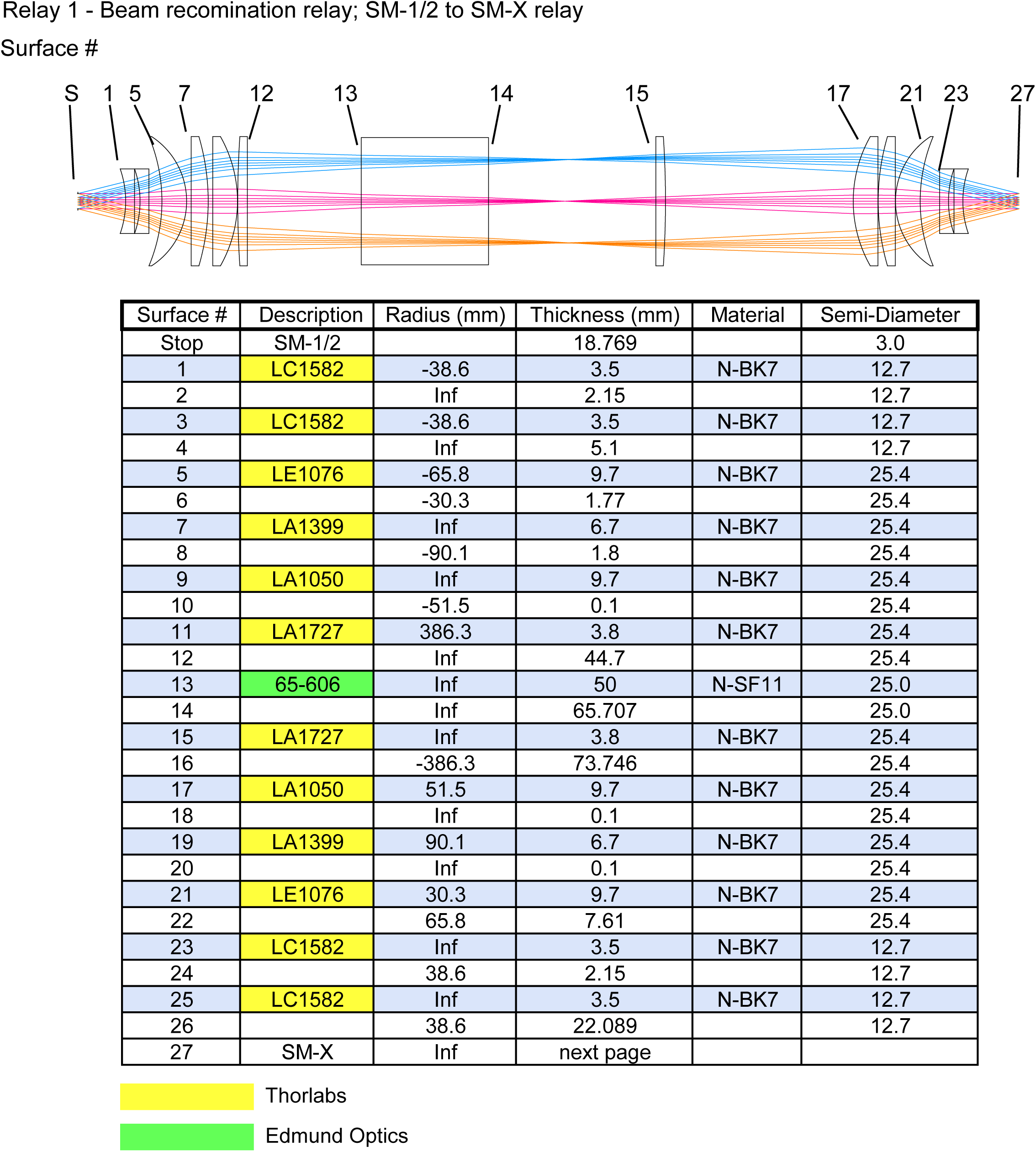
Full prescription data for the steering mirror (SM) to X scanning mirror (SM-X) relay. The optical relay was constructed from commercial off-the-shelf (COTS) components and was designed to minimize aberrations at high scan angles. The polarizing beam splitter (PBS) was offset from the focal point of the relay to minimize photo-damage to the optical cement.

**Supplementary Figure 2.**
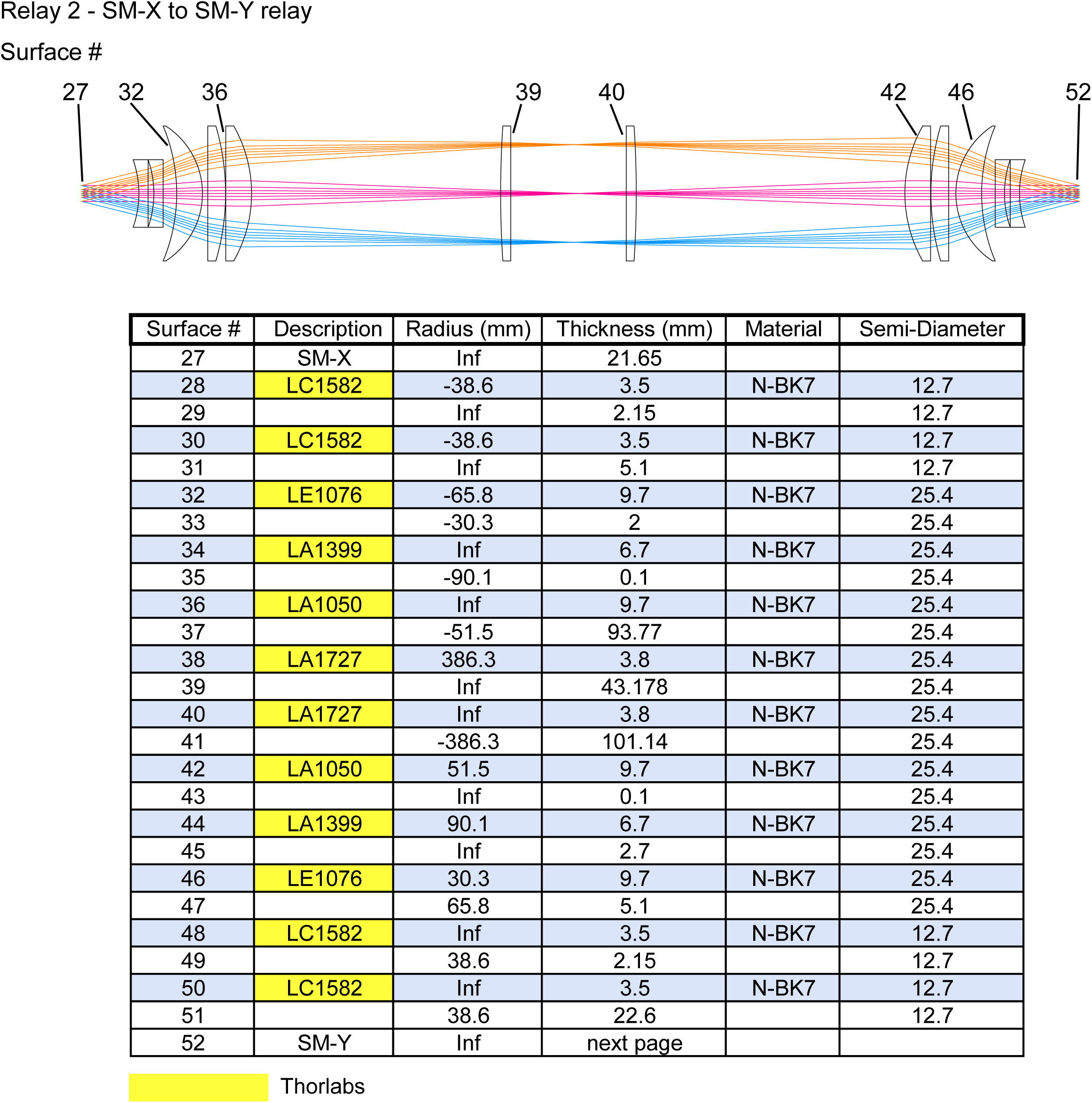
Full prescription data for the X scanning mirror (SM-X) to Y scanning mirror (SM-Y) relay. The optical relay was constructed from COTS components and was designed to minimize aberrations at high scan angles.

**Supplementary Figure 3.**
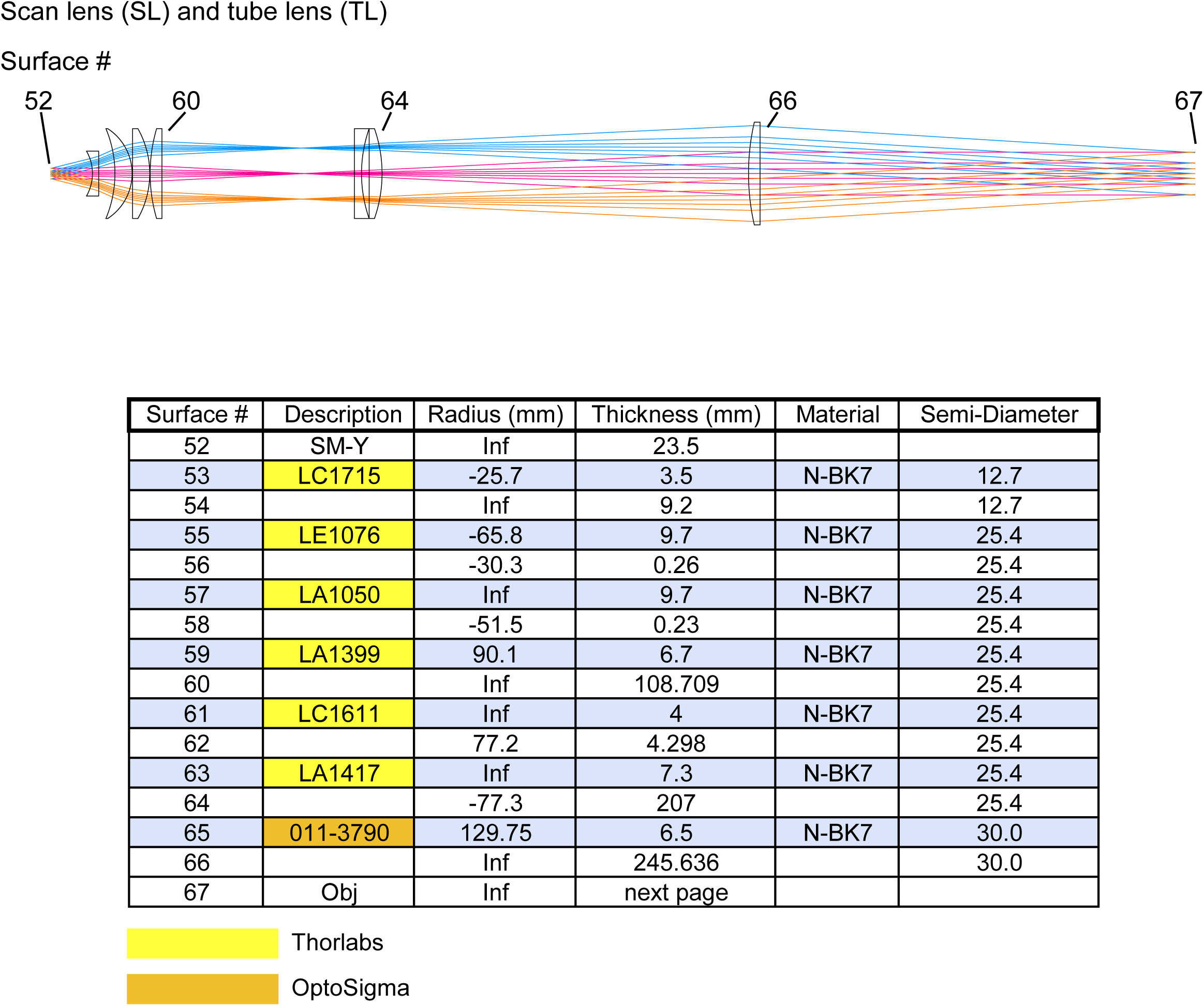
Full prescription data for the scan lens and tube lens optical subsystem. The scan and tube lens subsystem was constructed from COTS components. The terminal lens in this system has a 60 mm diameter in order to minimize vignetting at the extreme scan angles.

**Supplementary Figure 4.**
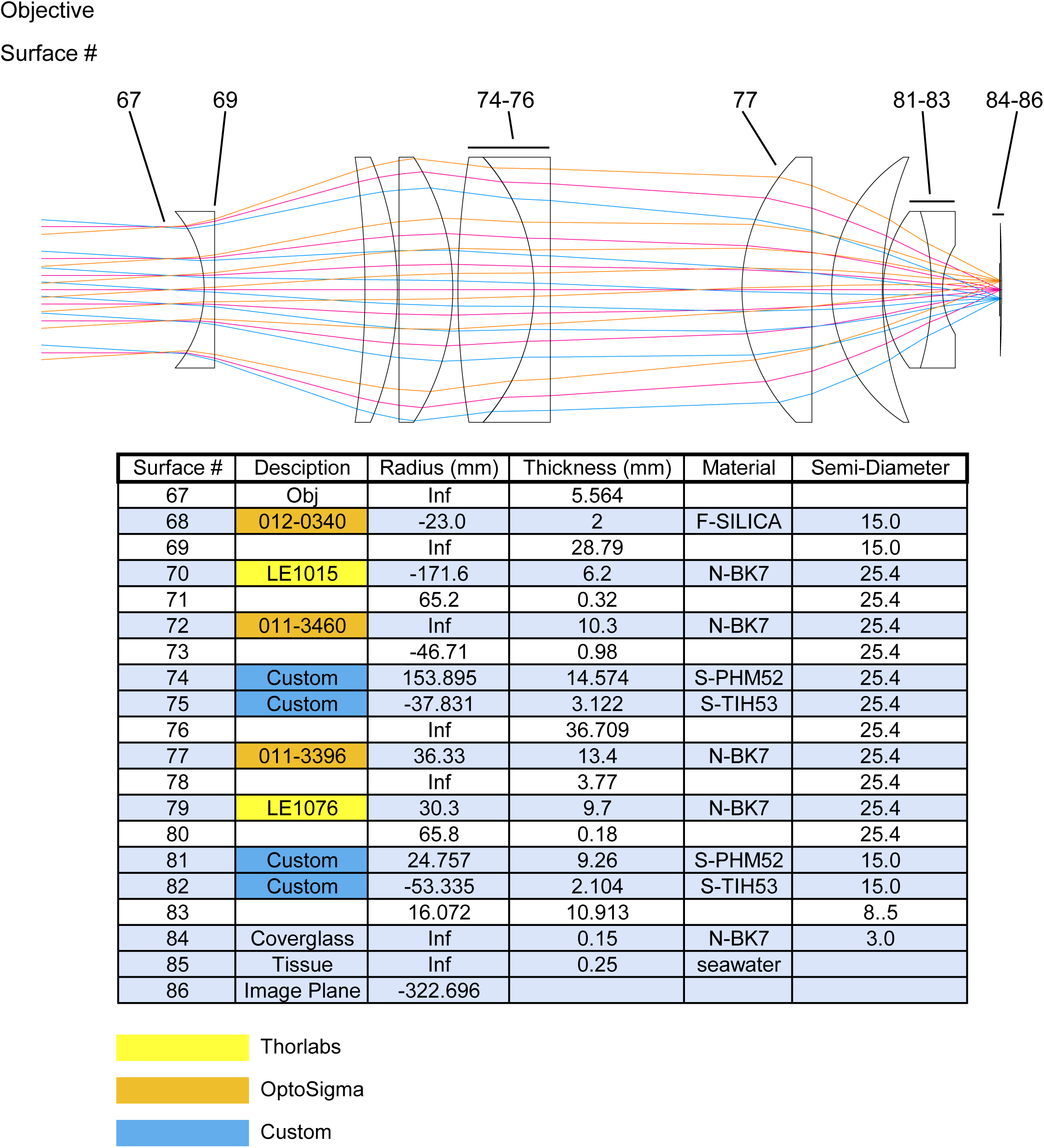
Full prescription data for the objective lens. The objective lens was constructed from COTS components and two custom cemented achromats. The objective has ∼8.5 mm of working distance.

**Supplementary Figure 5.**
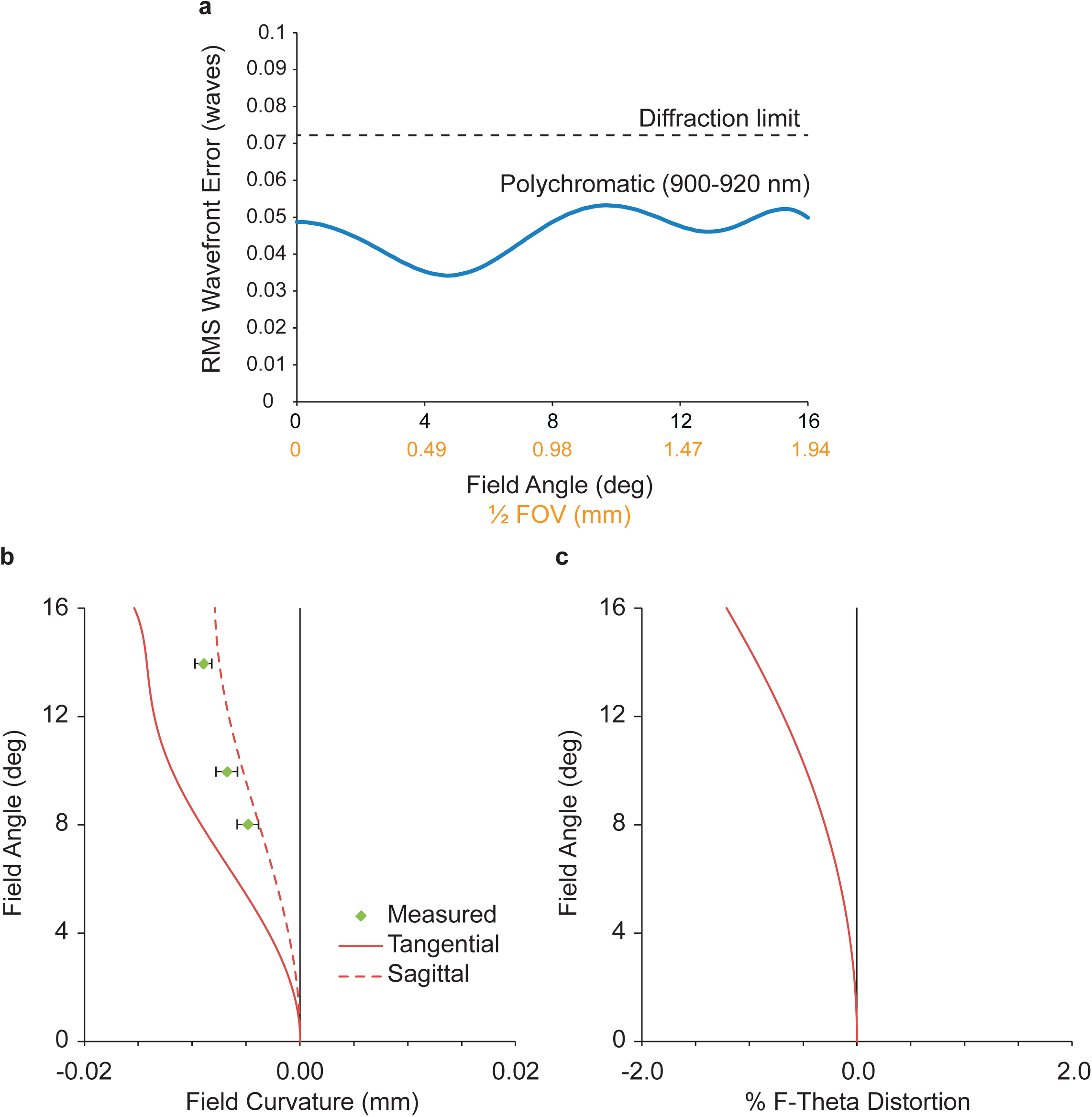
Simulations of the Trepan2p optical performance. **(a)** The Trepan2p system was optimized for diffraction limited performance across ∼4 mm FOV. **(b)** The simulated tangential and sagittal field curvatures are less than 20 μm. The experimental field curvature is shown and was found by measuring the optimal *Z* focus of submicron beads. **(c)** F-theta distortions are minimal. The field curvature and F-theta distortion were sufficiently small to permit measurements of neural activity, and the minimal wavefront error was a major design criterion to ensure effective 2p excitation.

**Supplementary Figure 6.**
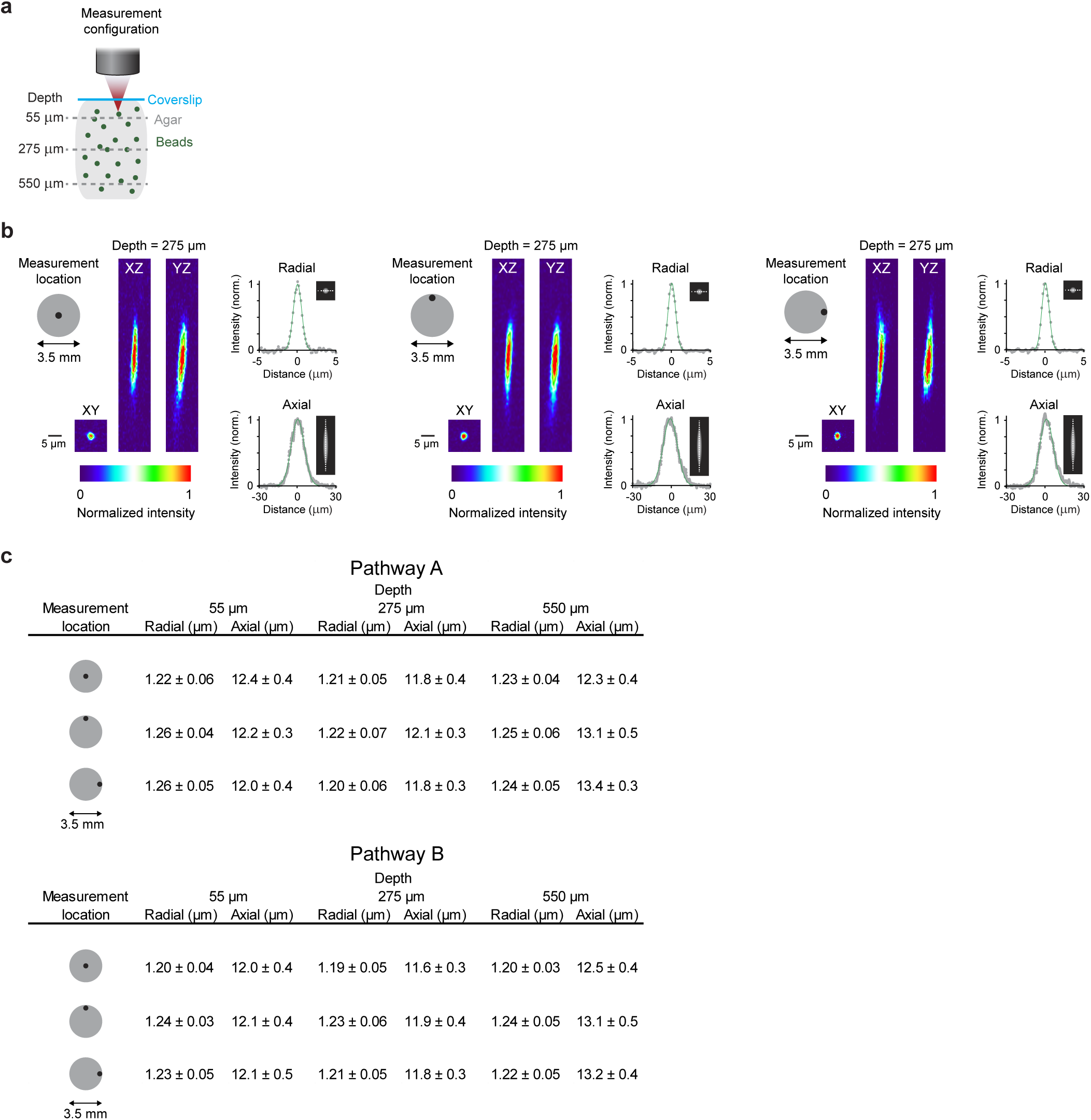
Focal excitation volume profile of the Trepan2p system. (**a**) 0.2 μm fluorescent beads were embedded in 0.75% agarose gel. 50 μm z-stacks were acquired, each centered at one of three depths (55 μm, 275 μm, 550 μm). This was done on axis, and at the edges of the 3.5mm field of view. (**b**) Radial and axial excitation volume measurements were made at the indicated locations and depths by fitting a Gaussian curve to the intensity profiles of the beads in the XY plane (measured in both the X and Y directions and averaged) and in the Z direction (measured in Z in both the XZ and YZ planes and averaged). (**c**) A summary of the excitation volume measurements at three depths, for three locations, and for both of the temporally multiplexed beam pathways are shown (full width at half maximum of the Gaussian fits +/- the standard deviation for measurements from 8 different beads). The excitation volume typically increases in axial extent with imaging depth beyond the optimized focal plane (250 μm), but the optimized aberration correction and moderate NA combine to largely mitigate that effect and preserves the excitation volume across the full field and hundreds of microns of imaging depth.

**Supplementary Figure 7.**
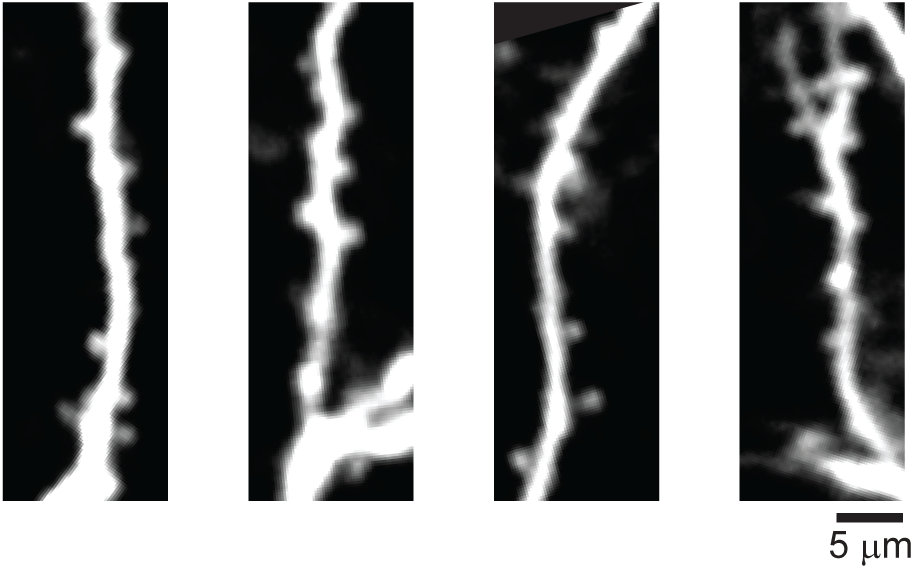
Spine imaging with the Trepan2p. Spines along the apical dendritic tufts of Layer V cells (Thy1-GFP, line O) are clearly visible at high magnification through the custom objective.

**Supplementary Figure 8.**
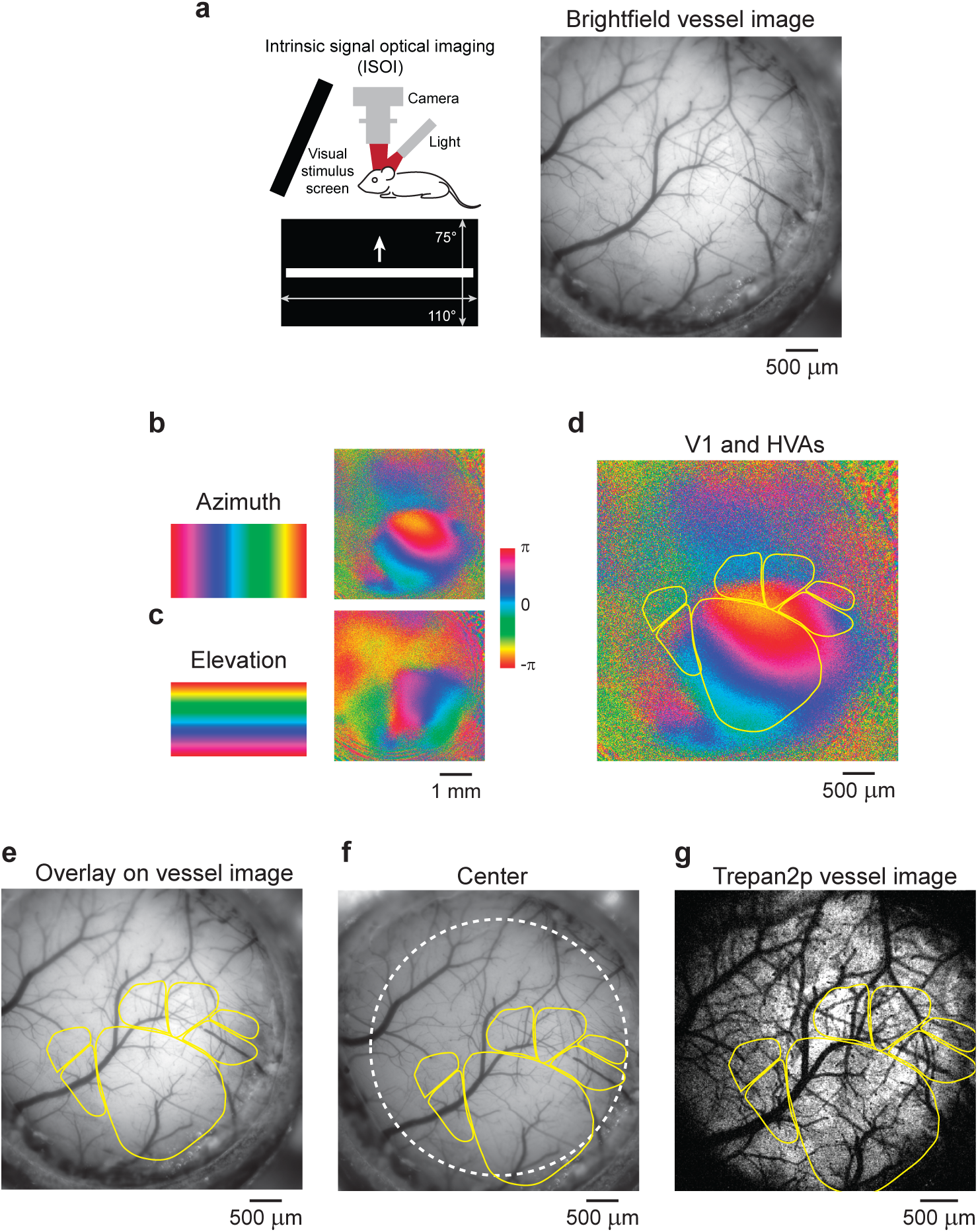
Identification of cortical areas across imaging modalities. Intrinsic signal optical imaging (ISOI) is a hemodynamic modality in which the reflectance of light off of the brain is modulated hemodynamic changes, which are in turn linked to stimulus-driven neural activity. ISOI can be used to map cortical activation at a resolution sufficient to delineate functional cortical area boundaries, and can be registered to maps of cortical vasculature (vessel image). (**a**) A vessel image was acquired prior to intrinsic signal optical imaging (ISOI) to map V1 and HVAs using retinotopic maps evoked by a single drifting (**b**) vertical (to map the azimuth of the cortical retinotopic maps in V1 and HVAs) or (**c**) horizontal bars (to map the elevation of the cortical retinotopic maps in V1 and HVAs). Phase of the evoked response is encoded in color and ranges from –π to +π, which corresponds to the two sides of the monitor (top and bottom when mapping elevation, or left and right when mapping azimuth). (**d**) These maps are used to locate the boundaries of V1 and HVAs. (**e**) These boundary lines were overlaid on the vessel map. The vasculature is imaged both in (**f**) the ISOI camera (the white line denotes the 3.5 mm FOV) and (**g**) the 2p imaging, and thus the vasculature is used to register between the two imaging modalities.

**Supplementary Figure 9.**
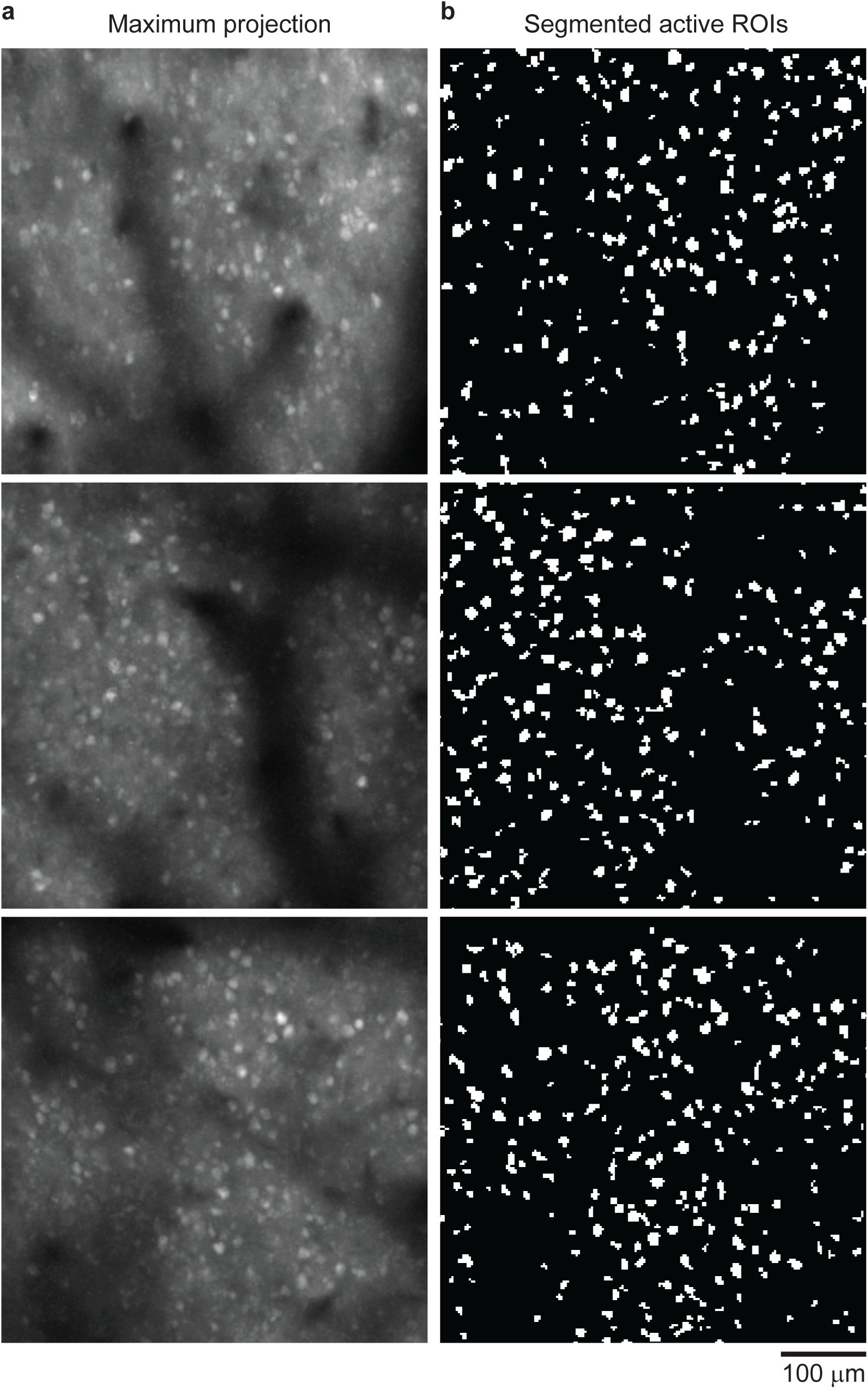
Magnified sub-regions from 3.5 mm field of view. (**a**) The maximum projections and (**b**) ROI maps inlayed in **Figure 1d** have been enlarged for visibility.

**Supplementary Figure 10.**
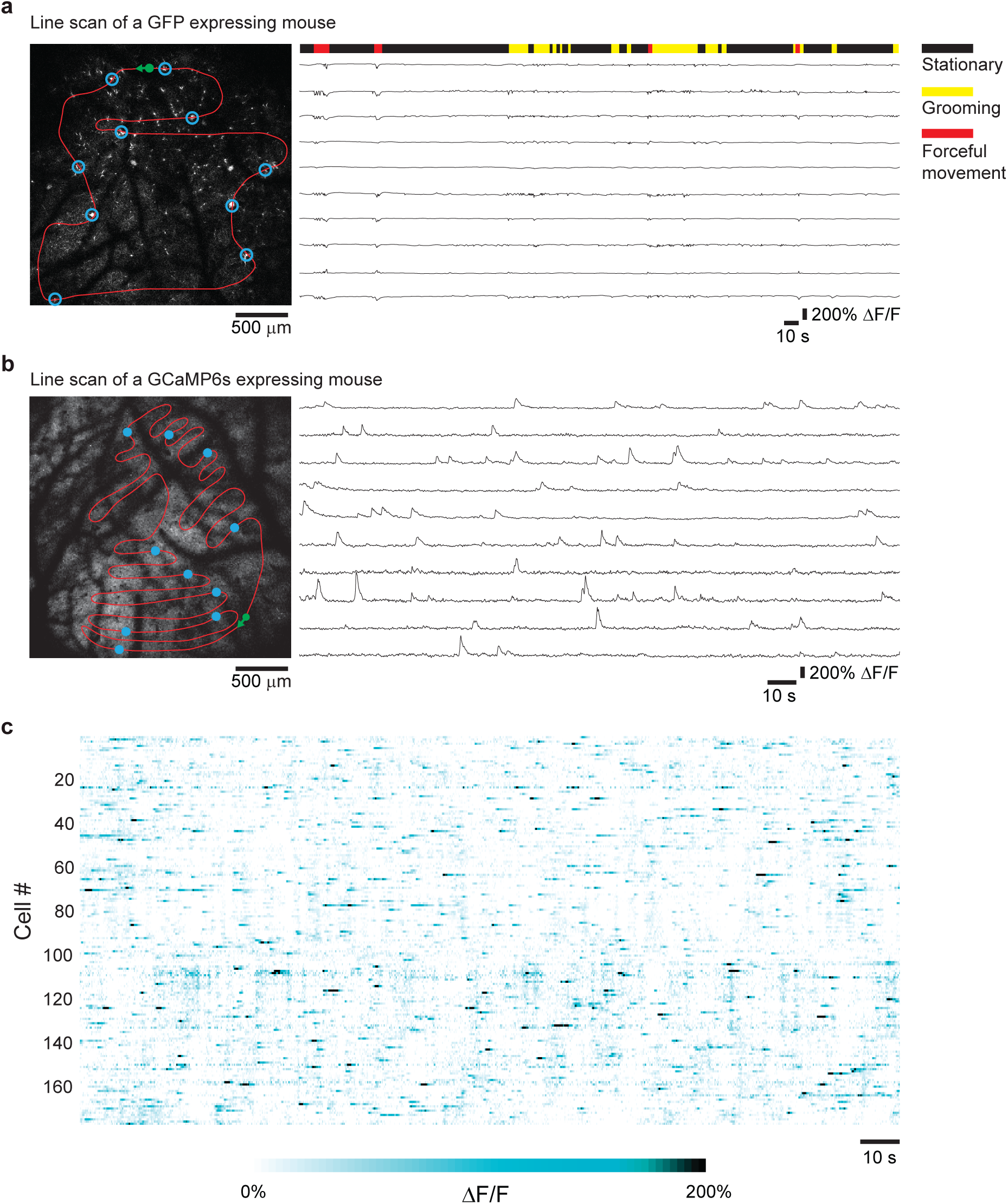
Fast imaging cycle times across a large field of view using arbitrary line scan in awake animals. (**a**) Line scan acquired at 96.1 Hz in an awake mouse expressing GFP in layer 5 neurons (Thy1-GFP, line O). The fluorescent structures in the left panel are ascending dendritic trunks from layer 5 neurons. Shown in red is the line scan path with the starting point and direction labeled in green. The open blue circles show the location of the traces shown in the right panel. The right panel is ΔF/F traces from ten structures in the line scan. Movement state of the mouse was tracked with simultaneous video monitoring and displayed above the traces. (**b**) Line scan acquired at 28.7 Hz in an awake mouse expressing GCaMP6s. Shown in red is the line scan path with the starting point and direction labeled in green. The closed blue circles show the location of the traces shown in the right panel. The right panel are ΔF/F traces from ten cells in the line scan. (**c**) Fluorescent transients (ΔF/F) from 172 active neurons along the scan path in (**b**). Note that the amplitude of GCaMP6 signals is typically larger than the movement-induced transients in panel (**a**).

**Supplementary Figure 11.**
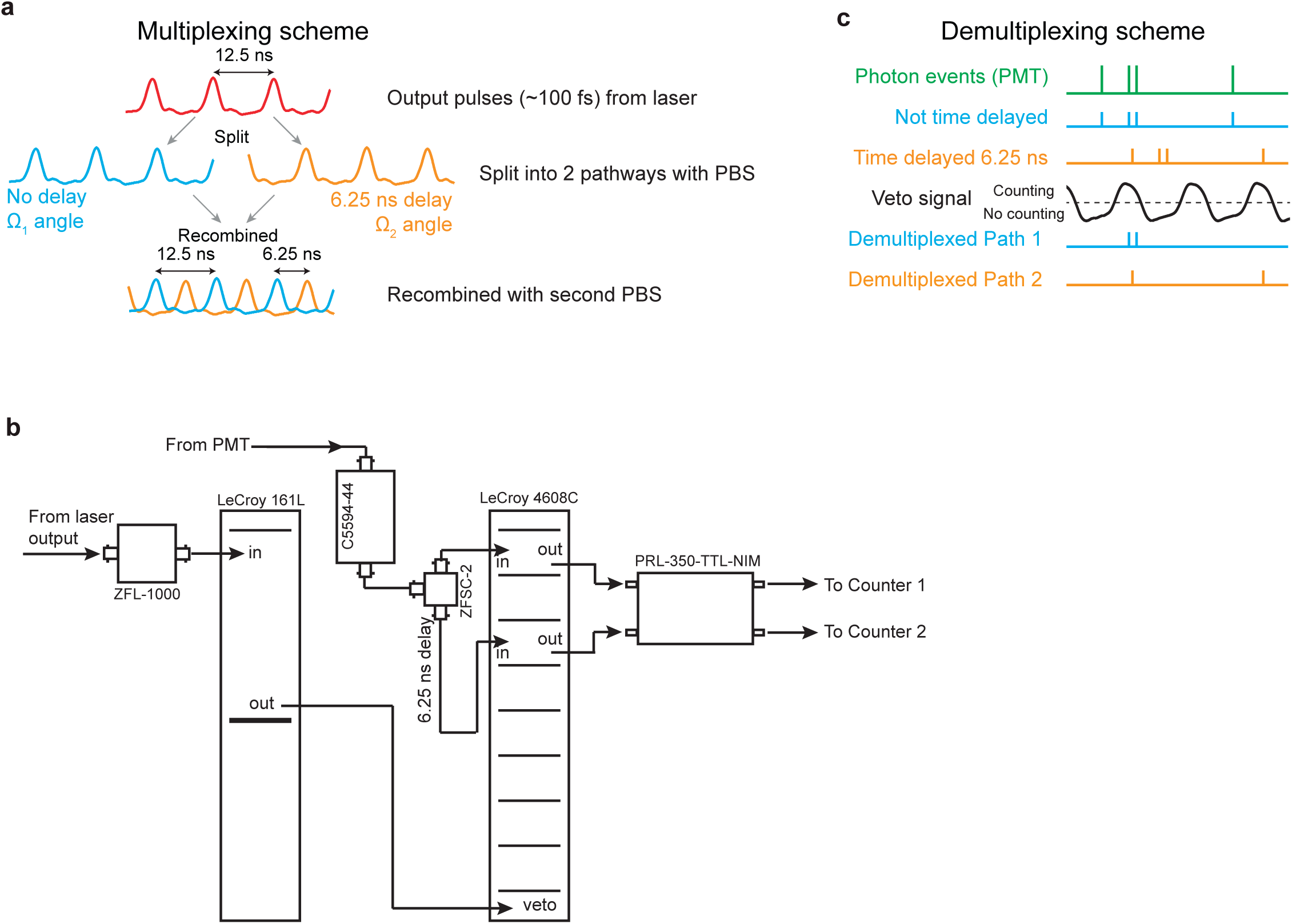
Multiplexing and demultiplexing. (**a**) In the multiplexing scheme, laser pulses are split into two pathways, and pathway 2 is delayed. The 6.25 ns delay in pathway 2 results in perfectly interleaved pulses upon recombination. Both pathways have different deflection angles (Ω_1_ or Ω_2_) on the galvanometer scanning mirrors, as dictated by SM_1_ and SM_2_. (**b**) High bandwidth electronics are used for demultiplexing and photon counting. (**c**) PMT pulses are equally divided into two pathways. The second pathway is time delayed by 6.25 ns. A veto pulse width of 6.25 ns is applied to the two detection pathways resulting in two demultiplexed output pulse streams. The veto signal prevents counting, and is active during the low state (*i.e.,* when the veto signal is high, counting is allowed). These events are counted and assembled into images.

**Supplementary Figure 12.**
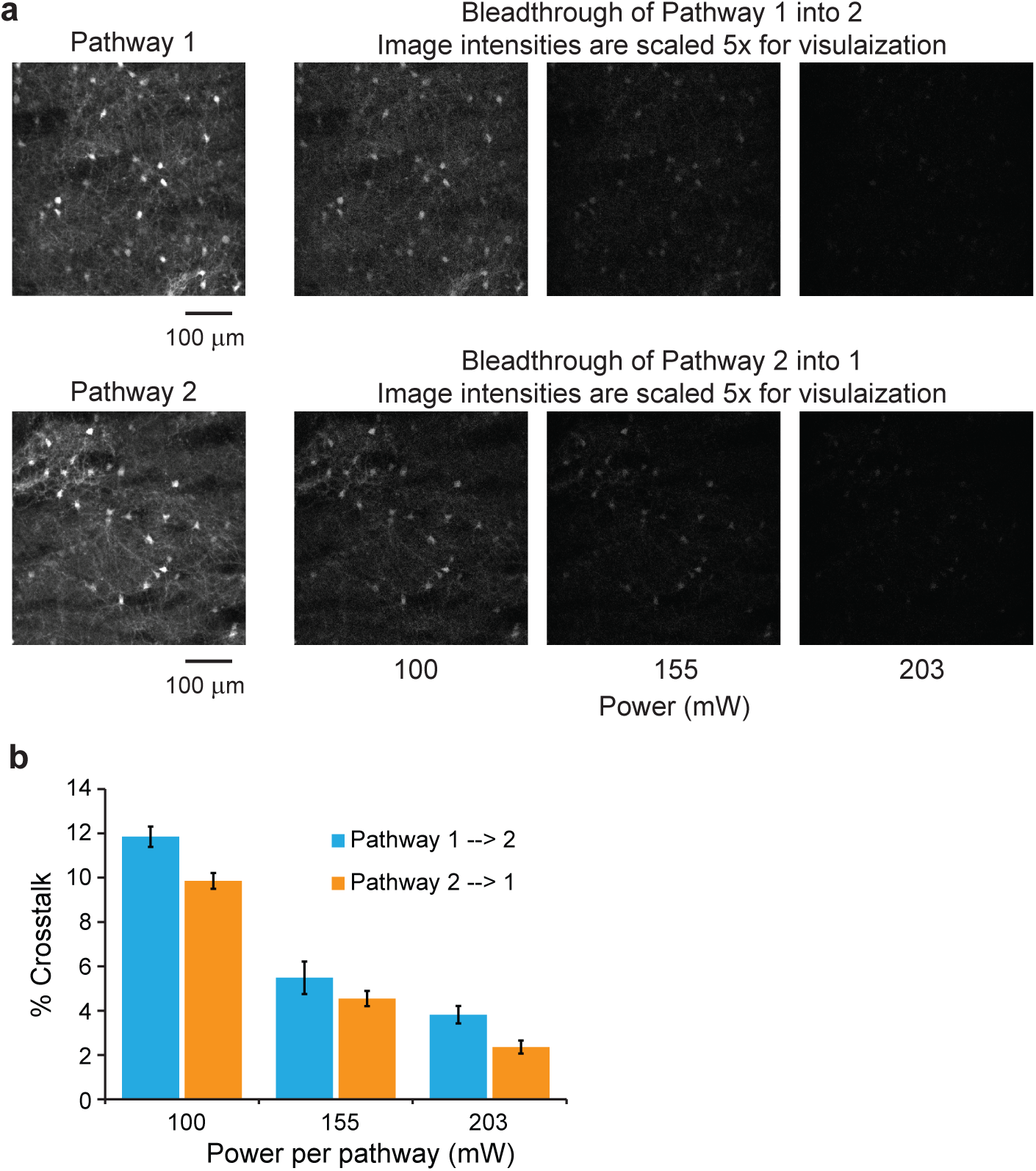
Crosstalk is minimal between the two pathways. (**a**) When measuring the crosstalk between the multiplexed pathways, the dynamic changes in the fluorescence intensity of GCaMP6s complicates analysis. Thus, we imaged neurons expressing YFP in an acute brain slice. Neurons were (top) imaged with pathway 1 while blocking excitation in pathway 2, and (bottom) vice versa. Any signal measured in the blocked pathway is due to crosstalk from the unblocked pathway. Crosstalk was measured with increasing levels of input power. (**b**) Mean fluorescent intensity of individual ROIs (somas) between the two pathways was measured, and decreased for increasing laser power due (see **Online Methods**). For the *in vivo* imaging described in this study, we used laser power of ∼195 mW, which indicates < 5% crosstalk between pathways.

**Supplementary Figure 13.**
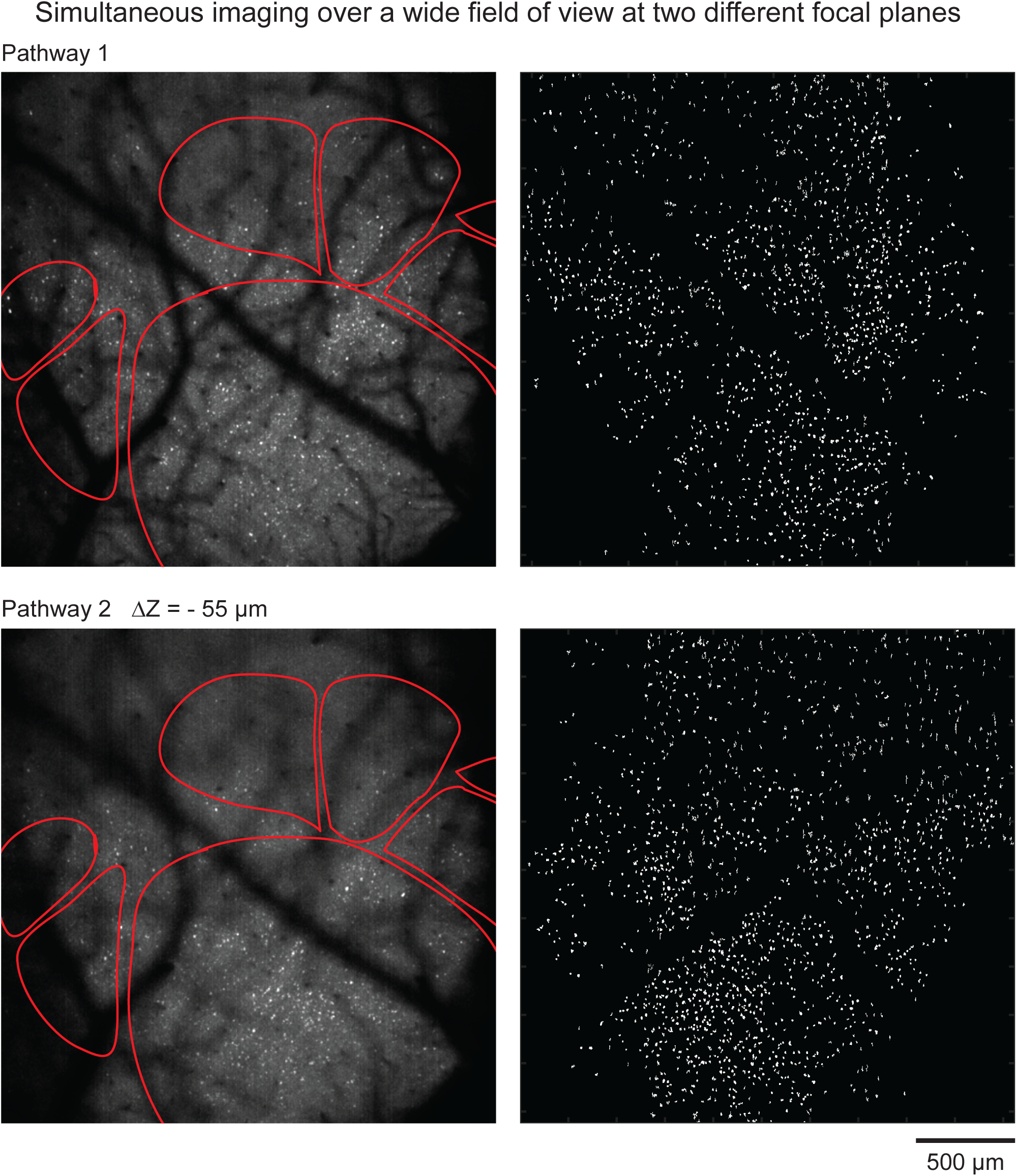
Dual depth imaging across a 2.5 mm field of view. The majority of the primary and higher visual areas are contained in a 2.5 mm field of view. Both imaging pathways were utilized to image (0.8 frames/s) the field of view at different focal planes (ΔZ = −55 μm). Left: maximum projection of the time series; right: binary image showing active ROIs. Data is shown in **Supplementary Video 7**.

**Supplementary Figure 14.**
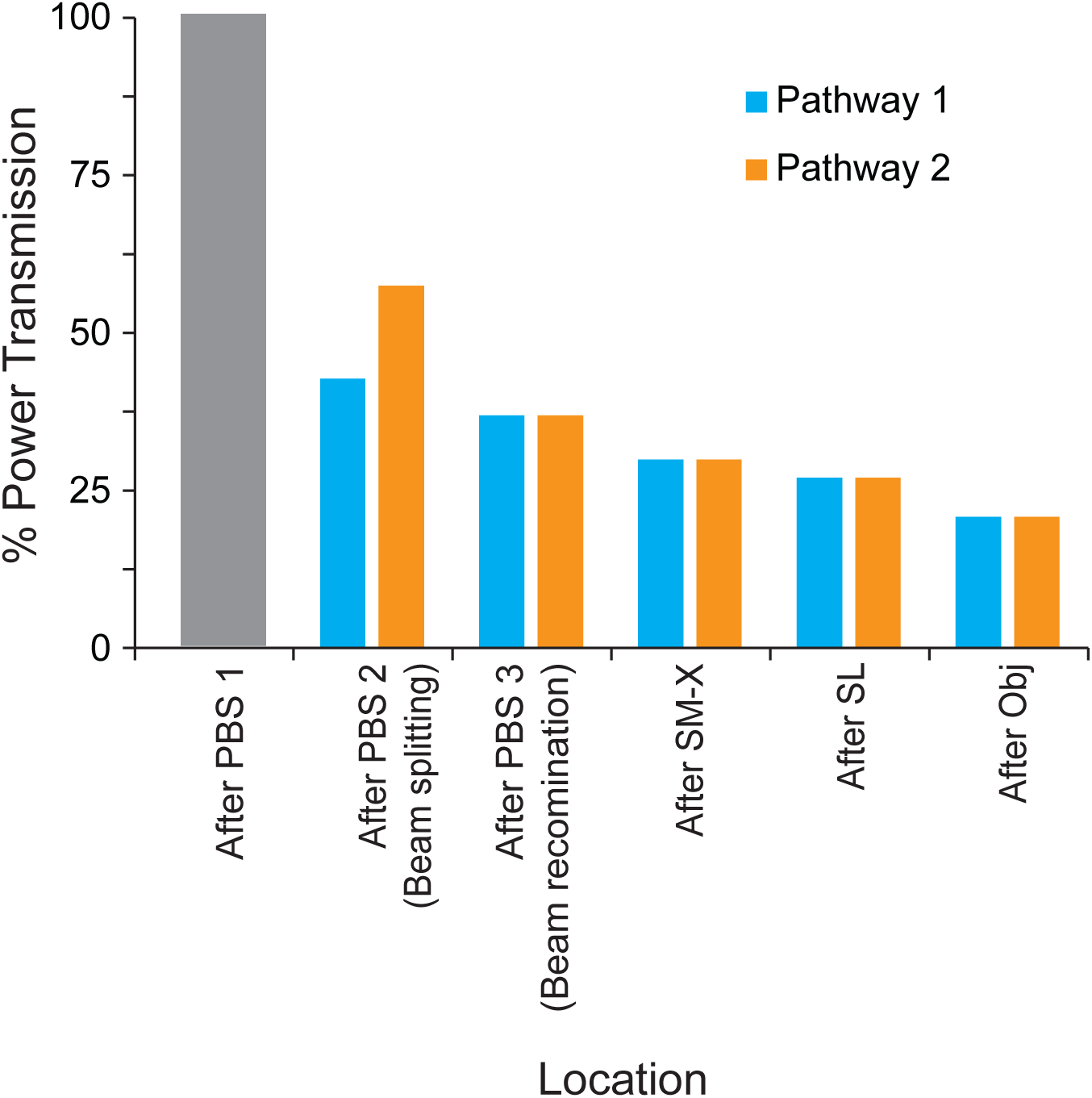
Power transmission through the Trepan2p. After attenuating the overall power using the first polarizing beam splitting cube, the power at subsequent locations in the Trepan2p microscope was measured. The second half-wave plate was adjusted such that the final power out of the front of the objective was identical between the two pathways. The larger losses in pathway 2 are due to the additional expansion (and subsequent clipping) the beam undergoes as it travels the additional 1.87 m path length. Overall, the system transmits 41% (20.5% per path) of the initial power.

## Supplementary Videos

**Supplementary Video 1. High resolution imaging using the Trepan2p.**

Neuronal activity in a mouse expressing GCaMP6s in neocortical pyramidal neurons was imaged through a cranial window while a naturalistic visual stimulus was presented. Images were acquired at 1024 × 1024 pixels at 1.9 frames/s. Video playback is 15 frames/s (∼7x real time).

**Supplementary Video 2. Z stack of GFP expressing layer 5 neurons.**

A Z-stack acquired in a Thy1-GFP O-line mouse *in vivo*. Layer V somas are readily visible >600 μm below the pial surface. The laser power was not adjusted during data acquisition. Intensity of the images was adjusted after acquisition for ease of visualization of the somas.

**Supplementary Video 3. Imaging neural activity across a 3.5 mm FOV.**

Neuronal activity in a mouse expressing GCaMP6s in neocortical pyramidal neurons was imaged through a cranial window while a naturalistic visual stimulus was presented. Images were acquired at 2048 × 2048 pixels at ∼0.1 frames/s. Video playback is 12 frames/s <u>(∼120x real time)</u>. The anatomical location imaged is shown on the right. Zooming in shows single neuron resolution is maintained across the FOV.

**Supplementary Video 4. Simultaneous dual region imaging at the same XY position and different depths.**

Neuronal activity in a mouse expressing GCaMP6s in neocortical pyramidal neurons was imaged through a cranial window while a naturalistic visual stimulus was presented. Images were acquired at 512 × 256 pixels at ∼9.5 frames/s. Video playback is 30 frames/s <u>(∼3x real time)</u>. The two imaging pathways (Pathway 1: left; Pathway 2: right) are located at the same XY location within V1 (**Fig. 2d**) and differ by 48.4 μm in Z

**Supplementary Video 5. Quad region imaging.**

Neuronal activity in a mouse expressing GCaMP6s in neocortical pyramidal neurons was imaged through a cranial window while a naturalistic visual stimulus was presented. Four regions were imaged by utilizing the two imaging pathways (Pathway 1: top row; Pathway 2: bottom row) and serially applying an offset voltage to the galvanometer scanners. Each of the four regions was acquired at 512 × 512 pixels at ∼1.9 frames/s. Video playback is 15 frames/s (∼7x real time).

**Supplementary Video 6. Quad region imaging, 1x real time playback.**

Same video as **Supplementary Video 5** with the playback in real time.

**Supplementary Video 7. Imaging neural activity across a 2.5 mm FOV simultaneously at two depths.**

Neuronal activity in a mouse expressing GCaMP6s in neocortical pyramidal neurons was imaged through a cranial window while a naturalistic visual stimulus was presented. The two imaging pathways (Pathway 1: left; Pathway 2: right) are located at the same XY location and differ by 55 μm in Z. Images were acquired at 2048 × 1024 pixels at 0.8 frames/s. Video playback is 12 frames/s (∼15x real time).

## ONLINE METHODS

### Animals

All procedures involving living animals were carried out in accordance with the guidelines and regulations of the US Department of Health and Human Services and approved by the Institutional Animal Care and Use Committee at University of North Carolina. C57Bl/6 mice were housed under a reversed 12hr-light/12hr-dark light cycle with ad libitum access to food and water. Two lines of transgenic mice were used that express GCaMP6s in cortical pyramidal neurons: (1) Cre-dependent GCaMP6s mice, Ai96 (RCL-GCaMP6s), crossed with Emx1-Cre mice (Jackson Labs stock #005628; **Fig. 1**) and (2) Cre/Tet-dependent GCaMP6s mice, Ai94 (TITL-GCaMP6s) triple crossed with Emx1-Cre and ROSA:LNL:tTA mice (Jackson Labs stock #011008; **Figs. 2, 3**). Ai94 and Ai96 lines were kindly provided by H. Zeng and the Allen Institute. We used a transgenic mouse in which a tyrosine hydroxylase (TH) promoter directs expression of Cre recombinase (TH-Cre; (Jackson Labs stock #008601). In this mouse, the ventral tegmental area (VTA) was injected with adeno-associated virus particles (AAV5-DIO-ChR2(H134R)-EYFP; 500 nL bilateral at an infusion rate of 100 nL/min). The animal was sacrificed three months post-surgery. TH-expressing terminals arising from the VTA were imaged in the striatum (**Supplementary Fig. 12**). Additionally, a Thy1-GFP O-line mouse, which expresses GFP largely in Layer V, was used to generate **Supplementary Figure 10** and **Supplementary Video 2**.

#### Surgery

Mice were deeply anesthetized using isoflurane (5% for induction, 1-2% for surgery) augmented with acepromazine (0 – 0.4 mg/kg body weight, i.p.). Physically-activated heat packs (SpaceGel, Braintree Scientific) were used to maintain the body temperature during surgery. The scalp overlaying right visual cortex was removed, and a custom head-fixing imaging chamber^22^ with 5-mm diameter opening was mounted to the skull with cyanoacrylate-based glue (Oasis Medical) and dental acrylic (Lang Dental). A 4-mm diameter craniotomy was performed over visual cortex. Carprofen (4.4mg/kg body weight, s.c.) was administered postoperatively to all mice that underwent recovery surgeries before returning to the home cage. Mice were mounted on a holder via the chamber^22^. For intrinsic signal optical imaging, this chamber was filled with a physiological saline containing (in mM): 150 NaCl, 2.5 KCl, 10 HEPES, 2 CaCl_2_ and 1 MgCl_2_.

### Intrinsic signal optical imaging (ISOI)

Custom instrumentation was adapted from the work of Kalatsky and Stryker^15^. Briefly, two F-mount lenses with respective focal lengths of 135 and 50 mm (Nikon) formed a tandem lens macroscope, which was attached to Dalsa 1M30 CCD camera (Teledyne DALSA). This configuration provided a 4.7 mm × 4.7 mm field of view (21.2 mm^2^). Acquired images were binned 2 × 2 spatially, resulting in a final pixel size of 9.2 μm × 9.2 μm. The pial vasculature was illuminated through a green filter (550 ± 50 nm, Edmund Optics) and the vasculature map was captured through a second green filter (560 ± 5 nm). From the pial surface, the macroscope was then focused down 600 μm where intrinsic signals were illuminated with halogen light (Asahi Spectra) delivered via light guides and focusing probes (Oriel) through a red filter (700 ± 38 nm, Chroma). Reflected light was captured through a second red filter (700 ± 5 nm, Edmund Optics) at the rate of 30 frames per second with custom-made image acquisition software (adapted by J.N.S. from code kindly provided by Dr. David Ferster, Northwestern University). Mice were head-fixed 20 cm from a flat 60 cm × 34 cm (width × height) monitor which was tilted towards the mouse 17.5° from vertical with their head angled to their right to cover the visual field (110° by 75°) of the contralateral eye. A light anesthetic plane was maintained with 0.5% isoflurane during imaging, augmented with acepromazine (1.5 - 3mg/kg), and the body temperature was kept at 37°C using feedback-controlled electric heat pad systems (custom-built).

### Visual stimuli

A drifting white bar on a black background (elevation and azimuth direction; 3° thick) was used to map retinotopy during ISOI. These were produced and presented using MATLAB and the Psychophysics Toolbox^23,24^. A corrective distortion was applied to compensate for the flatness of the monitor^25^ (code is available online, http://labrigger.com/blog/2012/03/06/mouse-visual-stim/). During calcium imaging, drifting gratings (0.04 cycles/°, 2 Hz, 8 directions, 10 s /direction) or a naturalistic movie were displayed on a small video display located 6.3 cm from the left eye. The naturalistic movie was from a helmet-mounted camera during a mountain biking run. This movie, while not something a mouse would typically encounter, provided a visual stimulus with a large amount of optic flow. Light from the display was shielded from the imaging apparatus using a shroud over the monitor.

### Image analysis for mapping cortical areas

Retinotopic maps obtained from intrinsic signal optical imaging were used to locate V1 and HVAs (**Supplementary Fig. 8**). Borders between these areas were drawn at the meridian of elevation and azimuth retinotopy. Magnitude response maps for grating patches were consulted for cross-verification. The vasculature acquired from Trepan2p imaging was compared to that acquired during ISOI and provided landmarks for identification of the locations within V1 and the HVAs.

### *In vivo* two photon imaging

All imaging was performed on the custom Trepan2p system. Control of the instrumentation (see **Trepan2p instrumentation** below) and image acquisition were controlled by custom LabVIEW software. Animals were lightly anesthetized (0.25% isoflurane) for the imaging in **Figure 1**; **Figure 2c, d**; **Supplementary Figures 7, 9**; and **Supplementary Videos 1-4**. Animals were awake for the imaging in **Figure 2e,f**; **Figure 3**; **Supplementary Figure 10, 13**; and **Supplementary Videos 5, 6**. The imaging was performed with ∼195 mW out of the front of the objective to minimize crosstalk (see **Crosstalk** below; **Supplementary Fig. 12**). Estimating the focal volume to be an ellipsoid with axes of 6 μm, 0.6 μm, and 0.6 μm (**Supplementary Fig. 6**), this equates to a power density of ∼21.5 mW/μm^3^. For a typical high NA objective (e.g. 16x, NA=0.8) the ellipsoid volume is ∼1.0 μm^3^ (axes of 1.5 μm, 0.4 μm, and 0.4 μm) and thus the power to have the equivalent power density would be 21.5 mW. The larger excitation volume allows us to increase the imaging power without the damage seen in higher NA objectives^26^. With typical imaging parameters (512×512 at 3.8 frames/s, 0.5 mm imaging region) up to 300 mW per channel could be used with no observed damage when imaging somas ∼ 150 microns deep. Damage was observed with these parameters if the imaging plane was moved to the surface of the dura. When imaging superficially, the power was kept below 100 mW.

### Image analysis for neuronal calcium signals

Ca^2+^ signals were analyzed using custom software in MATLAB (Mathworks). Neurons were identified and segmented using either a pixel-wise correlation map^6^ or a pixel-wise kurtosis map. For the kurtosis map, each imaging frame of the calcium imaging video was first Gaussian filtered in XY (sigma = 2 pixels), and then the kurtosis for the time series of each pixel location was computed using the filtered version of the video. This yields a single real-valued map with enhanced contrast for regions of interest that exhibit larger fluorescence transients. The maps were segmented into individual ROIs (neurons and processes) using a locally adaptive threshold and ΔF/F traces were calculated from the raw, unfiltered data. An exponential moving average of time width ∼150 ms was applied to the high speed ΔF/F traces in **Fig. 2e,f and Fig. 3a**, all other data was left unfiltered. For the correlation analysis, spike inference^6,27,28^ was performed on the raw ΔF/F traces prior to computing correlations. For pairwise correlations >0.5, traces were inspected for crosstalk. If the ROIs were present in the same XY positions in both imaging pathways and had high correlation values, the ROI with the weaker signal was determined to be due to crosstalk and was omitted (see **Crosstalk** below).

### Arbitrary line scan

The line scan pathway was drawn in custom LabVIEW software and converted to voltage commands for the galvanometer scanners. The imaging rate (28.7 Hz; **Supplementary Fig. 10b**) was determined to yield approximately equivalent pixel dwell time as the raster scans. Line scan data was arranged into a location-on-path versus time plot. The kurtosis of the data was taken along the time axis to yield a vector containing peaks (high values of kurtosis along the scan path). A small portion of the path (3-5 pixels) surrounding a peak of kurtosis was summed along the path length axis to yield a fluorescence time series for an active neuron. Each potential active region was manually inspected. This yielded 172 active cells distributed over the field of view (**Supplementary Fig. 10c**). An exponential moving average of time width ∼150ms was applied to the traces in **Supplementary Figure 10a, b**.

### Excitation volume measurements

Sub-micron fluorescent beads (0.2 μm, Invitrogen F-8811) were imbedded in a thick (∼1.2 mm) 0.75% agarose gel. 50 μm z-stacks were acquired, each centered at one of three depths (55 μm, 275 μm, 550 μm). This was done on axis and at the edges of the field of view for both imaging pathways. The minimal laser power for excitation was used to minimize the effects of non-linear expansion of the apparent excitation volume^29^. Gaussian curves were fit to the data to extract FWHM measurements. Data reported (**Supplementary Fig. 6**) are the mean ± standard deviation of 8 beads.

### Optical design and simulations

Relay, scan, tube, and objective lens systems were modeled in OpticStudio (Zemax, LLC). Three optical subsystems ([1] Relay. [2] Scan and Tube Lens, [3] Objective) were first designed, modeled, and optimized individually. After initial isolated design, the system was simulated and optimized as a whole to ensure there would be no unforeseen additive aberrations between subsystems. One of the main difficulties in designing a highly corrected optical system is dealing with chromatic aberrations. We decided to focus our design around a narrow wavelength range suitable for GCaMP excitation (910 ± 10 nm). This eased our requirements for correction of chromatic aberrations. For the relay and scan and tube systems, all commercial off-the-shelf (COTS) lenses (Thorlabs and OptoSigma) were used to minimize cost (**Supplementary Figs. 1-3**). These lenses were coated with an IR anti-reflective coating that had <0.5% reflectance at 910 nm.

To obtain diffraction limited performance (**Supplementary Fig. 5a**), the cost-effective COTS components were supplemented with two custom cemented doublets (**Supplementary Figs. 1-4)**. The custom cemented doublets were manufactured by Rainbow Research Optics, Inc. (Centennial, CO) and were coated with a broad-band anti-reflective coating (<1% 400-1100 nm). The other elements were COTS (Thorlabs and OptoSigma) and coated with a broad-band anti-reflective coating (<1% 525-925 nm; OptoSigma). We relaxed the requirement to have the focal plane be flat as slight deviations of the focal plane across the FOV would not affect our ability to capture neural dynamics. The resulting Trepan2p field curvature is <20 μm (**Supplementary Fig. 5b**). Additionally, the F-theta distortion is minor (<2%) (**Supplementary Fig. 5c**). After the individual sub-systems were designed and optimized, the system was optimized as a whole, while further refinement of spacing was made to allow for optomechanical limitations (lens housing, scanning mirrors). Complete lens data of the Trepan2p system is given in **Supplementary Figures 1-4**.

### Trepan2p instrumentation

Laser pulses from a Ti:Sapphire laser (Mai-Tai; Newport) with an automated pre-chirper unit (DeepSee; Newport) were attenuated using a half wave plate followed by a polarization beam splitting cube. Similar polarization optics were used to split the beam into two paths and control the relative power between the two paths. Prior to splitting, the beam was expanded using a 3x beam expander (Thorlabs). One beam travels directly to a custom motorized steering mirror, and the other beam is first diverted to a delay arm, and subsequently to a separately controlled steering mirror (**Fig. 2a**). The delay arm is designed to impart a 6.25 ns temporal offset to the pulses in one beam (1.875 m additional path length). Since the laser pulses are delivered at 12.5 ns intervals (80 MHz), they are evenly spaced in time at 160 MHz after the two beams are recombined (**Supplementary Fig. 11a**).

The steering mirror in each pathway imparts a solid angle (Ω_1_ or Ω_2_) deflection to the beam prior to recombination (**Fig. 2a**). It is this angle that determines the central locations (X_1_, Y_1_; X_2_, Y_2_) of individual imaging subareas within the larger FOV. The two beams are combined and relayed to the X-axis galvanometer scanner using the third polarization cube located in the middle of an afocal pupil relay (**Supplementary Fig. 1**). The 2-inch polarization beam combining cube (Edmund Optics) was positioned in the afocal relay such that the focal point of the relay was outside the cube. This was to avoid high intensity damage to the cemented surface within the cube. A similar afocal pupil relay (**Supplementary Fig. 2**) is present between the X-axis and Y-axis galvanometers (Cambridge Technologies; **Supplementary Note 3**). In this manner the two beams are simultaneously raster scanned (field size determined by scan amplitude), but with independently controlled spatial positions (determined by Ω_1_, Ω_2_). Immediately prior to the steering mirror in Pathways 1 and 2 are electrically tunable lenses (Optotune)^21^ which provides a z-range of ∼450 μm.

A scan lens and tube lens form a 4x telescope (**Supplementary Fig. 3**) and relayed the expanded beams to the back aperture of the custom objective (EFL = 27.5 mm; NA = 0.4; **Supplementary Fig. 4**). The entrance scan angles on the objective were ∼±3.7 degrees yielding our 3.5 mm FOV. Overall, the system transmitted 41% of entering laser power (**Supplementary Fig. 14**). The main factors influencing this figure were the galvanometer scanners and the objective, which were both overfilled. The system was not power limited in this configuration. Decreasing the overfilling of these elements can increase power transmission, though with a possible decrease in excitation efficiency (effective NA). Fluorescence was collected using a dichroic mirror, two lenses^30^, and a GaAsP photomultiplier tube (PMT; H10769PA-40, Hamamatsu).

### Photon counting electronics

Photon counting provides an increase in signal to noise over analog integration for dim signals^31^ which are common with *in vivo* two-photon calcium imaging data. Output from the photomultiplier tube (PMT; H10769PA-40, Hamamatsu) was first amplified with a high bandwidth amplifier (C5594-44, Hamamatsu) and then split into two channels (ZFSC-2-2A, Mini-Circuits) (**Supplementary Fig. 11b**). One channel was delayed relative to the other by 6.25 ns by using different lengths of BNC cable (∼ Δ122 cm) (**Supplementary Fig. 11b, c**). These were connected to a NIM discriminator (4608C octal discriminator, LeCroy). The ∼80 MHz synchronization output pulses from the laser was amplified (Mini-Circuits ZFL-1000) and then delivered to a second NIM discriminator (161L dual discriminator, LeCroy) (**Supplementary Fig. 11b**) which has a continuous potentiometer adjustment to adjust the output NIM pulse width from ∼5 ns to >150 ns. This output pulse was delivered to the common veto input on the LeCroy 4608C discriminator where the PMT outputs were collected. The veto width was adjusted by the potentiometer on the LeCroy 161L discriminator and the relative phase of the veto window was adjusted by time delaying the synchronization pulses from the laser module using small lengths of cables (∼0.5 ns resolution). The demultiplexed output pulses from the LeCroy 4608C were sent to a NIM-to-TTL converter (PRL-350-TTL-NIM, Pulse Research Labs) before going to the counter inputs on a fast counter (PCIe-6353, National Instruments) (**Supplementary Fig. 11b**). Counter measurements were organized into images in the custom LabVIEW software. Corrections for the sinusoidal scanning path were performed online in software^32^. In this manner we could demultiplex the single PMT output into two channels corresponding to the two excitation pathways (**Supplementary Fig. 11c**).

### Crosstalk

Many factors affect the crosstalk between the two pathways including GCaMP6 fluorescence lifetime, veto pulse width, veto pulse phase, and excitation power. The veto width was set to inhibit pulses arising from the off-target pathway while maximizing the signal from the desired pathway. The working value is ∼6.25 ns though small deviations from this made little difference. The phase of the veto (inhibit counting) signal relative to the PMT signal played a more significant role and was adjusted to give maximal signal in one channel arising from the non-delayed excitation path (Pathway 1) and maximal signal in the other channel arising from the 6.25 ns delayed excitation path (Pathway 2), while at the same time minimizing the signal from pathway 1 excitation when Pathway 2 was blocked and similarly for Pathway 2 into Pathway 1 (**Supplementary Fig. 12a**) With a fluorescence lifetime of ∼2.7 ns (single-exponential, ref. ^33^), ∼10% of the integrated curve would lie outside of a 6.25 ns time window, and could contribute to crosstalk. With increases in the laser power, the effective crosstalk was greatly diminished (**Supplementary Fig. 12a-c**) due to pileup distortion, which results in an apparently faster decaying effective fluorescence lifetime curve^34^. By adjusting the laser intensity and gain on the PMT, we can achieve smaller levels of crosstalk than predicted, while still obtaining high signal-to-noise measurements of fluorescence transients. For the experiments shown in this study, the laser power was ∼195 mW out of the front of the objective (also see ***In vivo* two photon imaging** section of **Methods**). For crosstalk to influence the acquired Ca^2+^ imaging and measured correlations there must be an ROI identified in the off-target pathway that is in the same XY position as a true ROI (neuron) in the on target pathway. False ROIs can occur, and these are ROIs that arise from the off-target pathway and are solely due to crosstalk. These can be identified by correlations in neural activity Ca^2+^ transients that satisfy two criteria: (1) have a high degree of correlation (>0.5), and (2) occur in the same XY locations in both images (pathway 1 and pathway 2). Thus, identification of the false ROIs is computationally simple, and they can be automatically removed from further analysis. Crosstalk was not detected in the data set displayed in **Figure 3**. A more subtle form of crosstalk can occur when two neurons, one in each pathway, are scanned simultaneously. To determine whether this form of crosstalk is affecting cross-correlation measurements, we searched the data set for pairs of neurons, one in each pathway, that overlap in XY position, and thus are simultaneously scanned and could contaminate each other’s signals. For these pairs, we computed the correlation between amount of overlap (the intersection of XY pixel locations between the two neurons’ ROIs) and activity cross-correlation. If activity signals from one neuron are contaminating the other, there should be a trend for more overlapped neurons to also exhibit higher cross-correlations. However, this relationship was not significant (*R*^*2*^ = 0.0094, *P* = 0.55, *N* = 40 pairs), and thus even when cells are simultaneously scanned, crosstalk is low and high fidelity measurements of neural activity are obtained.

